# First Detection of Epizootic Haemorrhagic Disease virus in the European Union, Italy-2022

**DOI:** 10.1101/2022.11.23.517495

**Authors:** Alessio Lorusso, Stefano Cappai, Federica Loi, Luigia Pinna, Angelo Ruiu, Giantonella Puggioni, Annalisa Guercio, Giuseppa Purpari, Domenico Vicari, Soufien Sghaier, Stephan Zientara, Massimo Spedicato, Salah Hammami, Thameur Ben Hassine, Ottavio Portanti, Emmanuel Breard, Corinne Sailleu, Massimo Ancora, Daria Di Sabatino, Daniela Morelli, Paolo Calistri, Giovanni Savini

**Affiliations:** Istituto Zooprofilattico Sperimentale dell’Abruzzo e del Molise, Teramo-Italy; Istituto Zooprofilattico Sperimentale della Sardegna, Sassari-Italy; Istituto Zooprofilattico Sperimentale della Sicilia, Palermo-Italy; Institut de la Recherche Vétérinaire de Tunisie, Tunis, Tunisia; UMR Virologie, INRAE, Ecole Nationale Vétérinaire d’Alfort (ENVA), ANSES, Laboratoire de Santé Animale, WOAH-EHDV Reference Laboratory, Maison Alfort 94700, France; Service de Microbiologie, Immunologie et Pathologie Générale, École Nationale de Médecine Vétérinaire de Sidi Thabet, IRESA, Universitè de la Manouba, Tunisia; Direction Générale des Services Vétérinaires, Commissariat Régional au Développement Agricole de Nabeul, Nabeul, Tunisia

## Abstract

We describe the first detection in the European Union of the epizootic haemorrhagic disease virus (EHDV). EHDV-8 has been detected in cattle farms in Sardinia and Sicily. The virus has a direct Northern African origin as its genome is identical (>99.9% nucleotide sequence identity) to EHDV-8 strains detected in Tunisia in 2021/2022.

## Introduction

The epizootic haemorrhagic disease (EHD) is a WOAH-listed disease of wild and domestic ruminants caused by EHD virus (EHDV). EHDV infection in deer, particularly among white-tailed deer (*Odocoileus virginianus*) in North America, can cause high levels of mortality. EHDV is related to the bluetongue virus (BTV), etiological agent of the bluetongue disease of ruminants (BT). Both viruses belong to the genus *Orbivirus* (family *Sedereoviridae*) and circulate in multiple serotypes (1,2). Their viral genome comprises 10 linear segments (S1-S10) of double-strand RNA and the structural outer capsid protein (coded by S2) determines serotype specificity. Both viruses cause similar clinical signs in cattle and are transmitted by several species of biting midges of the genus *Culicoides*. BT primarily affects sheep and has been described multiple times in the last twenty-four years in the European Union (EU) causing devastating outbreaks in ruminants with repercussions on animal trade (3). Most European BT outbreaks had a direct Northern African origin because of wind-driven dissemination of BTV-infected midges from these areas (1, 4–7).

Here, we describe the detection of EHDV-8 in cattle in Sardinia and Sicily, Italy. This represents the first evidence of EHD in the European Union (EU). This report follows our previous studies on EHDV-8 in Tunisian cattle in 2021 (8).

## Materials and Methods

On October 28 2022, clinical signs suggestive of BT infection (inappetence, cyanosis and edema of the tongue, conjunctivitis, and fever) were reported by the Local Veterinary Services (LVS) in one cattle from a farm (F) 1 (Fig. 1A) located in the municipality of Arbus (Fig. 1B). On November 3, the animal succumbed. At necropsy, spleen was collected by the LVS along with blood samples from three additional symptomatic cattle. On November 4 2022, in F2 and F3 (Arbus and Guspini, respectively Fig. 1B) three cattle showed the same symptomatology (Fig. 1C). On October 25 2022, BT-like clinical signs were also evidenced in three cattle belonging to F4 located nearby Trapani (Sicily, Fig. 1D). Samples were collected from the described animals and tested for the presence of EHDV RNA (VetMAX^™^ EHDV Kit rRT-PCREHDV; Thermo Scientific™ Waltham, MA, USA). A real time RT-PCR specific for the S2 of EHDV-8 TUN 2021 (rRT-PCR_EHDV-8_) was established (Portanti et al., manuscript in preparation) as the available test designed on the S2 of the EHDV-8 reference serotype (Australia, 1982) was not able to detect EHDV-8 TUN 2021 (8). Overall, nucleotide sequences of forward and reverse primers and probes are EHDV_Ser8varNEW_fwd AGAGATGAAGATCGCGAGGA (975-994); EHDV_Ser8varNEW_rev GAATCACACGCGCTCACTAA (1135-1159) and EHDV_Ser8varNEW_Probe FAM-ACGGATGAGATACGGAACATACGGGG-TAMRA (1066-1091), respectively. Whole genome sequencing-WGS (9) was performed on selected samples to get information upon the genome constellation of the occurring strain.

**Fig 1.**
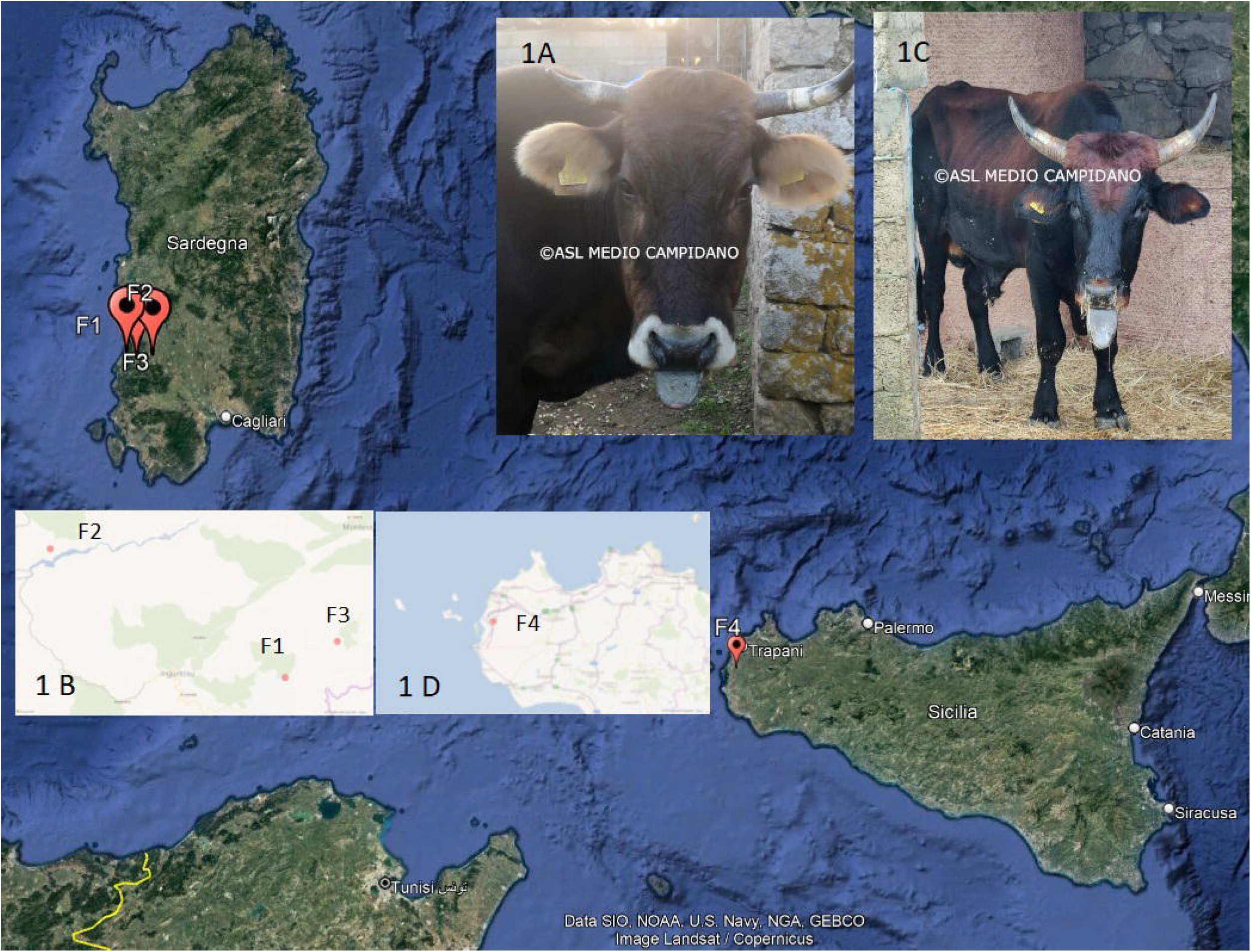
Geographical locations (Google Earth) of the four farms involved in the early outbreaks and clinical signs observed in cattle (cyanosis and edema of the tongue).

## Results

All sampled animals from Sardinia and Sicily were positive for EHDV (Ct range 23-28). Genotyping by a specific S2 EHDV-8 TUN2021 real-time RT-PCR confirmed the presence of a EHDV-8 TUN2021-like strain. The evidence of EHDV was notified to the Italian Ministry of Health which in turn notified the WOAH and the European Commission. This implied the ban of animal (all ruminants) movement from the two islands and the establishment of a 150 km radius restriction zone around the outbreaks. One EHDV-8 positive blood sample from Sardinia was selected for WGS. Genome analysis confirmed that the Sardinian EHDV-8 strain (EHDV-8 SAR2022 NCBI Submission ID: 2646388) shares the same genome constellation and high nucleotide sequence relatedness (>99.9%) with multiple EHDV-8 TUN 2021-like strains sequenced so far. WGS for Sicilian strains is currently ongoing.

## Discussion

A novel *Orbivirus* incursion to the EU sustained by EHDV-8 is reported. This virus has a direct Northern African origin. This event was reasonably predictable as for the current widespread distribution of the same virus in Tunisia and likely in neighboring countries (8), and in consideration of the previous incursions of multiple BTV strains to Southern Europe. At the time this report has been prepared, on November 18 2022, EHD has been notified also in Andalusia (Cadiz and Sevilla)-Spain. At this point, it is hard to predict the future scenarios for the EU cattle production system and the impact of the possible spread of EHDV. EHD will probably pose new challenges that the EU veterinary authorities will be forced to face. The lessons learned with BT should be a reference for choosing proper control and prevention strategies for EHD. Overall, these events further emphasize the importance for European countries in general, and for Italy due to its geographical location, of having in place robust collaborations with Northern African authorities on public and animal health. The prompt detection of EHDV-8 in Sardinia and Sicily is, indeed, the last example of the benefits that could derive from such relationships. This was crucial, as it facilitated the development of a specific and accurate molecular test for the detection of EHDV-8 as knowledge upon the genome constellation and the genomic relatedness of EHDV-8 with extant EHDV serotypes was already achieved. Undoubtedly, vaccine development needs to be boosted as vaccination is the only strategy to prevent direct and indirect economic losses and significantly reduce virus circulation.

## Ethical approval

No ethical approval was required as biological samples from animals were collected during outbreak investigations.

## Acknowledgements

The mention of commercial products in this article is simply for the purpose of providing specific information and does not imply a recommendation or approval by IZS-Te.

## References

1. Maclachlan NJ, Mayo CE, Daniels PW, Savini G, Zientara S, Gibbs EP. Bluetongue Rev Sci Tech. 2015; 34, 32

2. Savini G, Afonso A, Mellor P, Aradaib I, Yadin H, Sanaa M, Wilson W, Monaco F, Domingo M. Epizootic heamorragic disease. Res Vet Sci. 2011;91(1):1–17.

3. Rushton J, Lyons N. Economic impact of Bluetongue: a review of the effects on production. Vet Ital. 2015;51(4):401–6.

4. Lorusso A, Guercio A, Purpari G, Cammà C, Calistri P, D’Alterio N, Hammami S, Sghaier S, Savini G. Bluetongue virus serotype 3 in Western Sicily November 2017. Vet Ital. 2017; 53(4):273–275.

5. Lorusso A, Sghaier S, Di Domenico M, Barbria ME, Zaccaria G, Megdich A, Portanti O, Seliman IB, Spedicato M, Pizzurro F, Carmine I, Teodori L, Mahjoub M, Mangone I, Leone A, Hammami S, Marcacci M, Savini G. Analysis of bluetongue serotype 3 spread in Tunisia and discovery of a novel strain related to the bluetongue virus isolated from a commercial sheep pox vaccine. Infect Genet Evol. 2018; Apr; 59:63–71.

6. Cappai S, Rolesu S, Loi F, Liciardi M, Leone A, Marcacci M, Teodori L, Mangone I, Sghaier S, Portanti O, Savini G, Lorusso A. Western Bluetongue virus serotype 3 in Sardinia, diagnosis and characterization. Transbound Emerg Dis. 2019;66(3):1426–1431.

7. Calistri P, Giovannini A, Conte A, Nannini D, Santucci U, Patta C, Rolesu S, Caporale V. Bluetongue in Italy: Part I. Vet Ital. 2004; 40(3):243–51

8. Sghaier S, Sailleau C, Marcacci M, Thabet S, Curini V, Ben Hassine T, Teodori L, Portanti O, Hammami S, Jurisic L, Spedicato M, Postic L, Gazani I, Ben Osman R, Zientara S, Breard E, Calistri P, Richt JA, Holmes EC, Savini G, Di Giallonardo F, Lorusso A. Epizootic Haemorrhagic Disease Virus Serotype 8 in Tunisia, 2021. Preprints 2022, 2022110195 (doi: 10.20944/preprints202211.0195.v1).

9. Marcacci M, De Luca E, Zaccaria G, Di Tommaso M, Mangone I, Aste G, Savini G, Boari A, Lorusso A. Genome characterization of feline morbillivirus from Italy. J Virol Methods. 2016; 234:160–3.

